# Multitasking Recurrent Networks Utilize Compositional Strategies for Control of Movement

**DOI:** 10.1101/2025.09.10.675375

**Authors:** John Lazzari, Shreya Saxena

## Abstract

The brain and body comprise a complex control system that can flexibly perform a diverse range of movements. Despite the high-dimensionality of the musculoskeletal system, both humans and other species are able to quickly adapt their existing repertoire of actions to novel settings. A strategy likely employed by the brain to accomplish such a feat is known as compositionality, or the ability to combine learned computational primitives to perform novel tasks. Previous works have demonstrated that recurrent neural networks (RNNs) are a useful tool to probe compositionality during diverse cognitive tasks. However, the attractor-based computations required for cognition are largely distinct from those required for the generation of movement, and it is unclear whether and how compositional structure extends to RNNs producing complex movements. To address this question, we train a multitasking RNN in feedback with a musculoskeletal arm model to perform ten distinct types of movements at various speeds and directions, using visual and proprioceptive feedback. The trained network expresses two complementary forms of composition: an algebraic organization that groups tasks by kinematic and rotational structure to enable the flexible creation of novel tasks, and a sequential strategy that stitches learned extension and retraction motifs to produce new compound movements. Across tasks, population activity occupied a shared, low-dimensional manifold, whereas activity across task epochs resides in orthogonal subspaces, indicating a principled separation of computations. Models that perform multitasking without the use of feedback or embodiment struggle to form compositional representations, suggesting these mechanisms constrain the network solution space during training. Finally, we demonstrate rapid transfer to held-out movements via simple input weight updates, as well as the generation of target trajectories from composite rule inputs, without altering recurrent dynamics, highlighting a biologically plausible route to within-manifold generalization. Our framework sheds light on how the brain might flexibly perform a diverse range of movements through the use of shared low-dimensional manifolds and compositional representations.

## 1 Introduction

In both cognitive and motor domains, animals can perform related but unfamiliar tasks with relatively little training. How does the brain achieve this remarkable flexibility? One widely proposed explanation is compositionality, the reuse and combination of previously learned computational primitives to perform new tasks. For instance, a professional dancer can learn new routines far more quickly than a novice. This is likely because the dancer already possesses foundational movement motifs that can be recombined in novel ways. The dancer can also retrieve and deploy the patterns needed for a wide variety of movements. This contrasts with most current control systems and deep learning models, which largely focus on narrow settings.

Several recent studies have experimentally explored the neural basis of compositionality in cognitive settings [1–7]. This work has inspired the study of compositional representations in recurrent neural networks (RNNs) through multitasking and continual learning [8–12]. Throughout this paper, we use “multitasking” in the sense established in this literature, meaning a single network trained to perform multiple distinct tasks, not necessarily simultaneously [8, 9]. RNNs trained on multiple cognitive tasks develop modular computational units that are composed in different ways across tasks [8, 9]. This insight has not yet been obtained for models that produce movement. It is unclear whether the same principles apply in a setting that generates continuous movement patterns by recruiting a musculoskeletal system, rather than the discrete motifs of working memory, decision making, or contextual inference tasks.

Compositional strategies during movement have been examined from different perspectives experimentally and theoretically [13–19]. Recent work suggests that primates reuse shared computations across tasks [20–23]. Such reuse has been demonstrated through shared low-dimensional manifolds across diverse wrist-based motor tasks in macaques [20], consistent with the manifold hypothesis that the coordinated activity of neurons at the population level, not single-neuron tuning, is the primary computation towards movement generation [24, 25]. Through this lens, network modes *define* the building blocks of a computation, and a shared manifold allows for *reuse* of such modes across tasks. At the same time, studies have shown that the motor cortex uses orthogonal subspaces to isolate computations, particularly between preparatory and movement epochs [26–28]. More explicit compositional representations of movement have also been reported. Motor cortical activity during a task requiring multiple sub-skills was found to be higher-dimensional than expected, with distinct sub-skills occupying distinct subspaces that may be combined compositionally [18], and premotor cortex has been shown to represent drawn shapes as discrete, generalizable symbols that can be composed into sequences [19]. Do networks trained on a variety of motor tasks adopt a similar structure to share and isolate computations? If so, does that structure support compositional representations of movement?

To model the motor cortex controlling the body, the joint use of RNNs and musculoskeletal models has been key for a diverse range of model organisms such as hydra, flies, rodents, and macaques [29–34]. Such embodied systems can reveal important structures in the networks trained to drive them [35]. Although these frameworks have been used to study the correspondence between trained networks and recorded neural trajectories, they have largely not been leveraged for understanding multitasking and compositionality. Notable studies considering embodied control during multitasking are [31, 36], where rat skeletal models were trained to perform multiple tasks using reinforcement learning. The work of [36] in particular was the first to analyze the representation of multitasking networks performing movement related tasks. However, these works largely focus on representational similarities across diverse behaviors, as opposed to the compositional strategies a network uses across tasks and whether these can be reused for transfer.

In this study, we consider two specific forms of compositionality. The first, algebraic compositionality, refers to how the network may perform individual movements as an additive function of parts of other movements. We say a task is built compositionally if multiple primitive computations, such as fixed points, transient dynamics, or distinct axes encoding task related variables, are combined and shared across tasks. The second form of compositionality is sequencing, in which sub-movements are combined one after another to produce a more complex movement [37]. We refer to a segment of the hand trajectory, such as a single reach or curved extension, as a sub-movement, and to the pattern of population activity and local dynamics that generates it as a motif; the two are analyzed separately. Whether network models develop these forms of compositionality through multitasking alone has not yet been studied.

Here, we develop a closed-loop multitasking RNN that controls a musculoskeletal arm to perform ten distinct types of movements, and ask whether compositional structure emerges through multitask training alone. The network is a feedback controller trained end to end: it receives task instructions and continuous visual and proprioceptive feedback and produces muscle commands, with no imposed decomposition into forward and inverse models. In the Discussion we relate the strategies it learns to optimal feedback control and internal-model theories of motor control.

The tasks include extensions and retractions of movement patterns such as straight reaches, curved reaches, and more complex patterns. Compositional representations can be built from these tasks both algebraically, by grouping kinematic and rotational similarities in a structured manner, and sequentially, through the ability to stitch sub-movements together. We find that low-dimensional, shared subspaces are consistently utilized during the movement phase of all tasks, while computations across epochs are organized in orthogonal subspaces. We then show that tasks are represented algebraically in network state space, and demonstrate that a task can be performed using motifs learned during other tasks. Additionally, we show that the motifs learned during training can be sequenced together without explicitly learning this relationship. Critically, embodiment and the associated feedback help the network develop such compositional representations, which we demonstrate through ablations. Finally, we show that the network is able to transfer its learned motifs to perform novel movements quickly, by only training the weights corresponding to a new static rule input. Overall, our framework provides a foundation for understanding the network structure and motifs that underlie motor compositionality.

## 2 Results

### 2.1 Successful multi-task kinematics achieved through training

We design a suite of tasks analogous to the center-out paradigm commonly used with primates in experimental settings (Fig. 1b; see Supplemental for task details). We first define the terms used throughout. A trial is a single instance of the network performing one task, that is, controlling the arm to produce one movement in a given direction at a given speed. A condition is a specific combination of direction and speed within a task. We also define extension movements, which are movements in which the arm extends outwards towards a target from a center position, as well as retraction movements, in which the arm retracts back to the center.

**Fig. 1.**
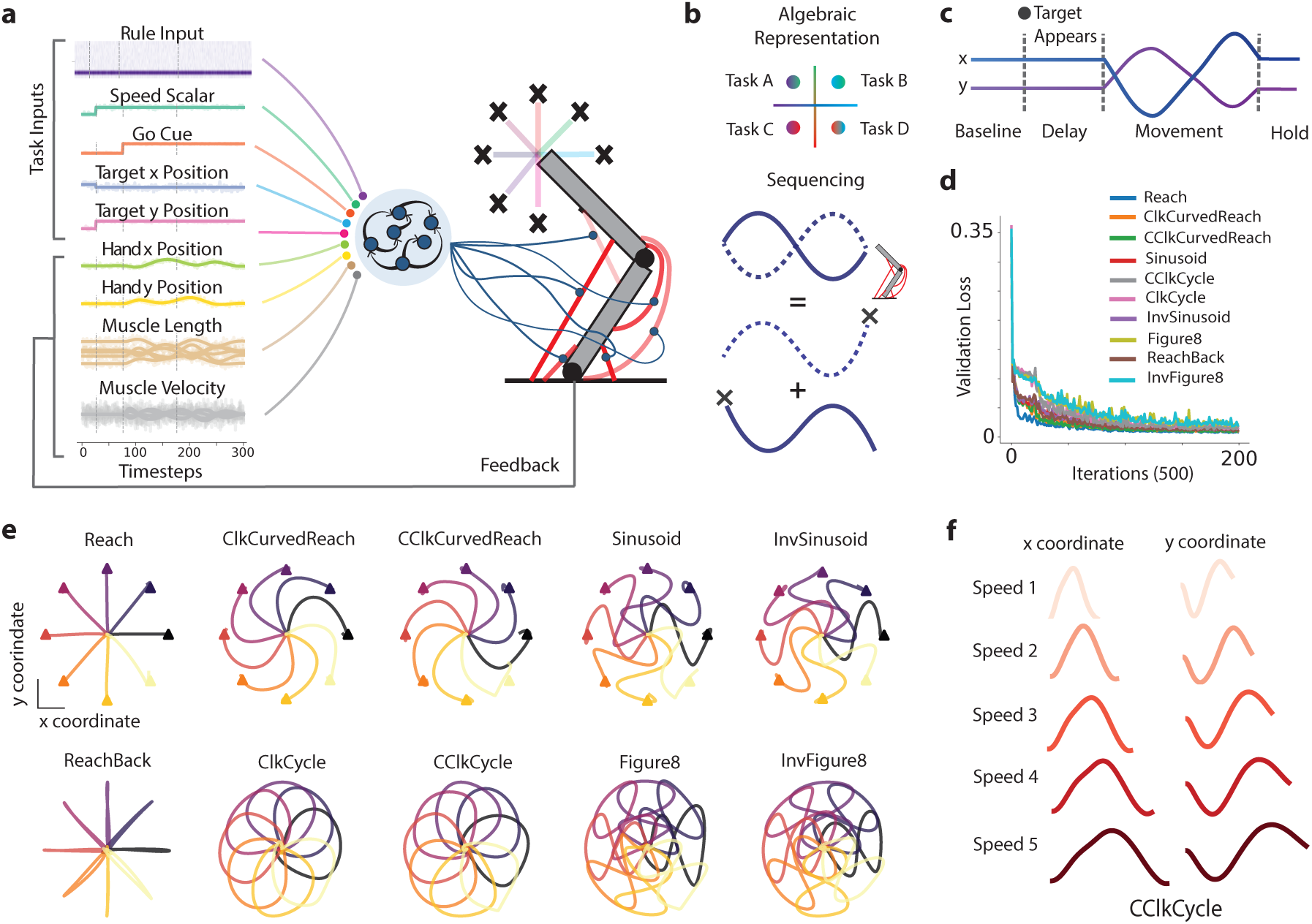
Task design and training performance. (a) We train an RNN in feedback with a musculoskeletal model in order to perform ten distinct kinematic trajectories in 16 different directions at five different speeds. Feedback from the arm is provided through muscle lengths, velocities, and visual feedback of the hand position. Other task-related variables are fed into the network such as the rule input, speed scalar, go cue, and target position. (b) Task design represents distinct kinematics containing composable elements. (top) Tasks can be constructed in an algebraic manner in network state space. (bottom) Individual extension and retraction motifs may be compartmentalized and sequenced in order to produce complex movements such as a figure eight. (c) Diagram of task structure. Task begins in a stabilization baseline period, where only the rule input appears, followed by a delay period in which condition-specific inputs appear, and finally movement occurs with a subsequent hold epoch. (d) Validation loss for interpolated conditions across all tasks. (e) Kinematic trajectories of the trained network for each task in each direction condition. (f) Example task kinematics for different speed conditions, with certain (not all ten) validation speeds included.

Each trial proceeds through four sequential epochs (Fig. 1c). In the baseline stabilization epoch, only the rule input is present and the arm settles to rest. In the delay epoch, condition-specific inputs (direction, speed, and target) appear, but the network must not yet move. In the movement epoch, the go cue activates and the arm executes the instructed trajectory. In the hold epoch, the arm maintains its final position. The delay epoch mirrors the preparatory period of delayed center-out reaching tasks in motor neuroscience [26, 38], and it lets us study preparatory activity before movement onset.

These tasks are intentionally simple. Well-defined motifs let us identify compositional structure cleanly and compare directly with established experimental paradigms. The tasks also share computations, which may allow the network to repurpose learned motifs. For example, the Figure8 task is composed of two sinusoidal movements, and its extension portion has the same kinematics as the Sinusoid task (Fig. 1b, bottom). Despite this kinematic similarity, these “subset” pairs (the Sinusoid is a subset of the Figure8) are distinguished by a one-hot contextual input that identifies the task, which we call the rule input. The curved reaches, cycles, sinusoids, and figure eights each define two tasks, one beginning clockwise and one counter-clockwise. If a compositional representation forms, the kinematic and rotational properties of the movements may be represented in a structured way that builds each one (Fig. 1b, top). Five tasks extend outward only, which we call extension tasks. Five tasks extend outward and then retract to the center using the same kinematic pattern, which we call extension-retraction tasks.

We trained an RNN with supervised learning to perform the ten motor tasks. Each task used 16 movement directions at five speeds, giving 80 conditions per task (Fig. 1a). For the arm musculoskeletal model, we use Motornet [29], a Python package for developing musculoskeletal models with differentiable muscles. The particular limb model we used is derived from [39], which implements a biologically plausible lumped-muscle model for 2D planar movements. The arm contains six muscles and two joints, and provides proprioceptive feedback to the network in the form of muscle lengths and velocities, and visual feedback of the hand position. The limb model we use here is simpler than those in previous embodiment studies [30, 31, 36], which presents a tractable framework for studying network representations and identifying motifs. Training produced accurate kinematic trajectories for all movements and conditions (Fig. 1e,f). Tasks were interleaved during training with equal probability. The validation loss dropped consistently for every task. Figure8 and InvFigure8 took longest to train and reached the highest final validation loss (Fig. 1d).

### 2.2 Within-task network dynamics are consistent with previous studies

Before analyzing structure across tasks, we first ask how the network solves each task on its own, and whether its solutions resemble those reported in the motor cortex. We use two standard tools. Principal component analysis (PCA) summarizes population activity in a few dominant modes, which shows how trajectories for different directions and speeds are arranged. jPCA detects rotational structure in the population trajectories, a signature commonly seen in motor cortex during reaching [38]. If these features match experimental observations, the model is a reasonable substrate for studying motor computation.

We begin with a single example task, InvSinusoid (counter-clockwise inverse sinusoid). During the delay epoch, the network settles onto a ring of fixed points, with one stable state per movement direction (Fig. 2b). These fixed points matter because they provide the initial conditions for movement. The network can hold a direction-specific preparatory state without moving, and the movement-epoch dynamics then unfold from that state. This is consistent with the initial-condition hypothesis of motor preparation [38, 40, 41].

**Fig. 2.**
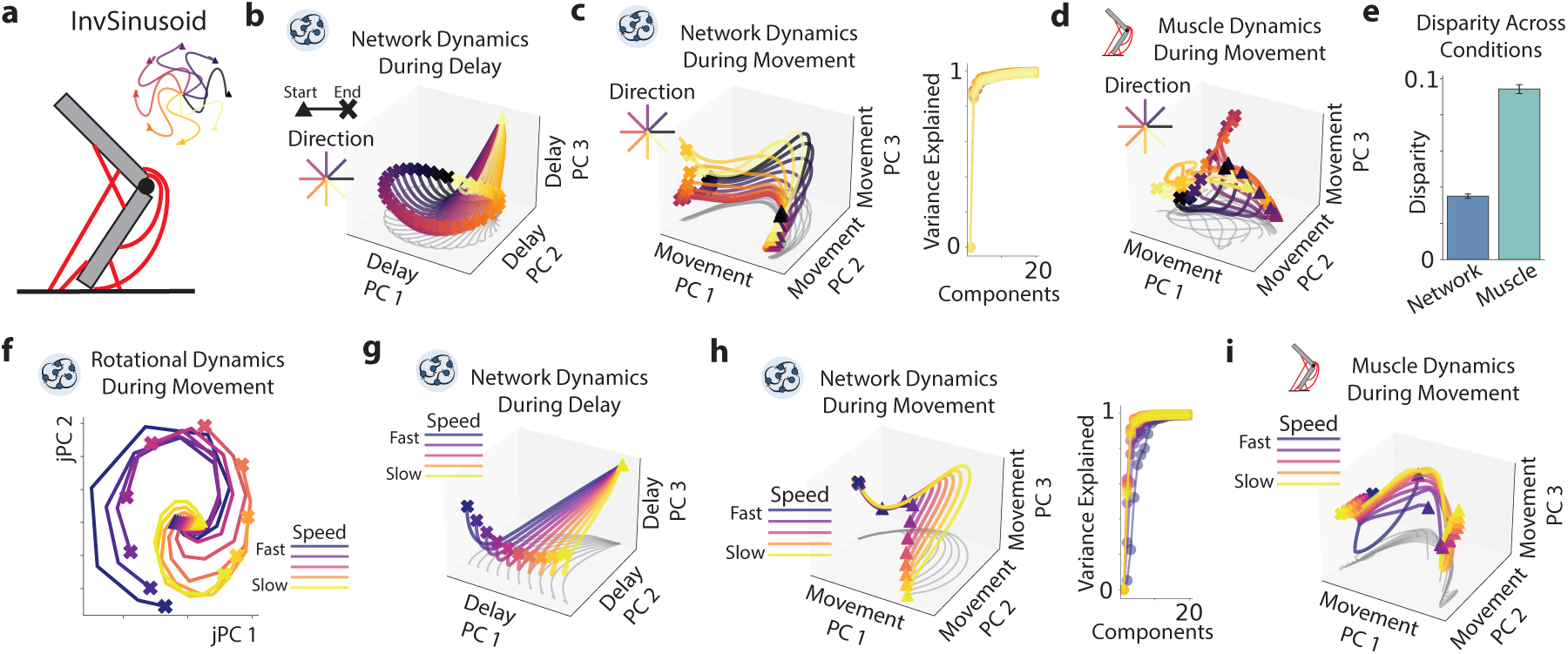
Within-task dynamics of trained network. (a) Example task, in this case an inverse sinusoid (InvSinusoid). (b) Network activity projected onto the first three PCs found during the delay epoch for the InvSinusoid task across 16 direction conditions. Network activity resembles a ring-like attractor predicting upcoming movement directions. Gray curves here and in later panels are 2D projections of the 3D trajectories onto the PC1-PC2 plane. (c) (left) Network activity projected onto the first three PCs found during the movement epoch across the same direction conditions. Network trajectories are offset according to the ring structure present during the delay. (right) Variance explained across directions demonstrate consistently low dimensional trajectories. (d) Same structure as (c) in the muscle space. Trajectories appear to be offset in a more complex fashion compared to network trajectories. (e) Procrustes similarity between conditions during the movement epoch for the network and muscles. Network uses more closely aligned trajectories than the muscles as shown in the PC plots. (f) jPCA computed across speed conditions shows that the network utilizes rotational dynamics. (g) Network activity projected onto delay PCs across speed conditions shows an offset along a single axis. (h) (left) Network trajectories projected onto the top three PCs shows speed conditions offset the same trajectories. (right) Variance explained across speed conditions shows consistent low dimensional trajectories. (i) Same as (h) for muscle activations.

Next, we show network trajectories trace similar shapes in the top PCs during the movement epoch across conditions, but they start from different points and are offset in magnitude along specific modes (Fig. 2c). We quantify this with a trajectory-shape similarity measure (procrustes analysis) across directions (Fig. 2e). For comparison, we show the muscle-space trajectories, which are less consistent (Fig. 2d). This pattern likely reflects the arm’s geometry, which requires slightly different solutions for different conditions. Other tasks show similar structure (Supplemental Fig. 2).

We next evaluate network activity across speed conditions. During the transient shift from delay to movement, the network engages in rotational dynamics (Fig. 2f), a feature commonly seen across motor cortex tasks [38]. During the delay and movement epochs, network trajectories appear as translated copies of the same trajectory with increasing rotation amplitude (Fig. 2g,h). This solution is similar but slightly weaker in the muscle space (Fig. 2i). The same dichotomy appears in M1 and muscle recordings from non-human primates [42]. Together, these results show that the model solves individual tasks using strategies similar to those reported in experimental and modeling studies. All other tasks use similar strategies as those shown here (Fig. 2).

### 2.3 Network shares neural modes and dynamics across similar movements

Having characterized how the network solves each task, we next ask how it organizes its activity across computations. We first consider the two epochs of a single task, preparation and movement, which serve different computational roles: holding a direction-specific state without moving, and then generating the movement itself. To compare the subspaces used in each epoch we use the variance-explained ratio, which asks how much of the activity in one epoch is captured by the principal components of the other, relative to its own components. A ratio near one means the two epochs use the same neural modes; a ratio near zero means their subspaces are orthogonal. Within every task, this ratio falls toward zero across the preparation and movement epochs (Fig. 3a), consistent with experimental findings in motor cortex [26, 27]. The network thus keeps distinct computations in distinct subspaces, even within a single task.

**Fig. 3.**
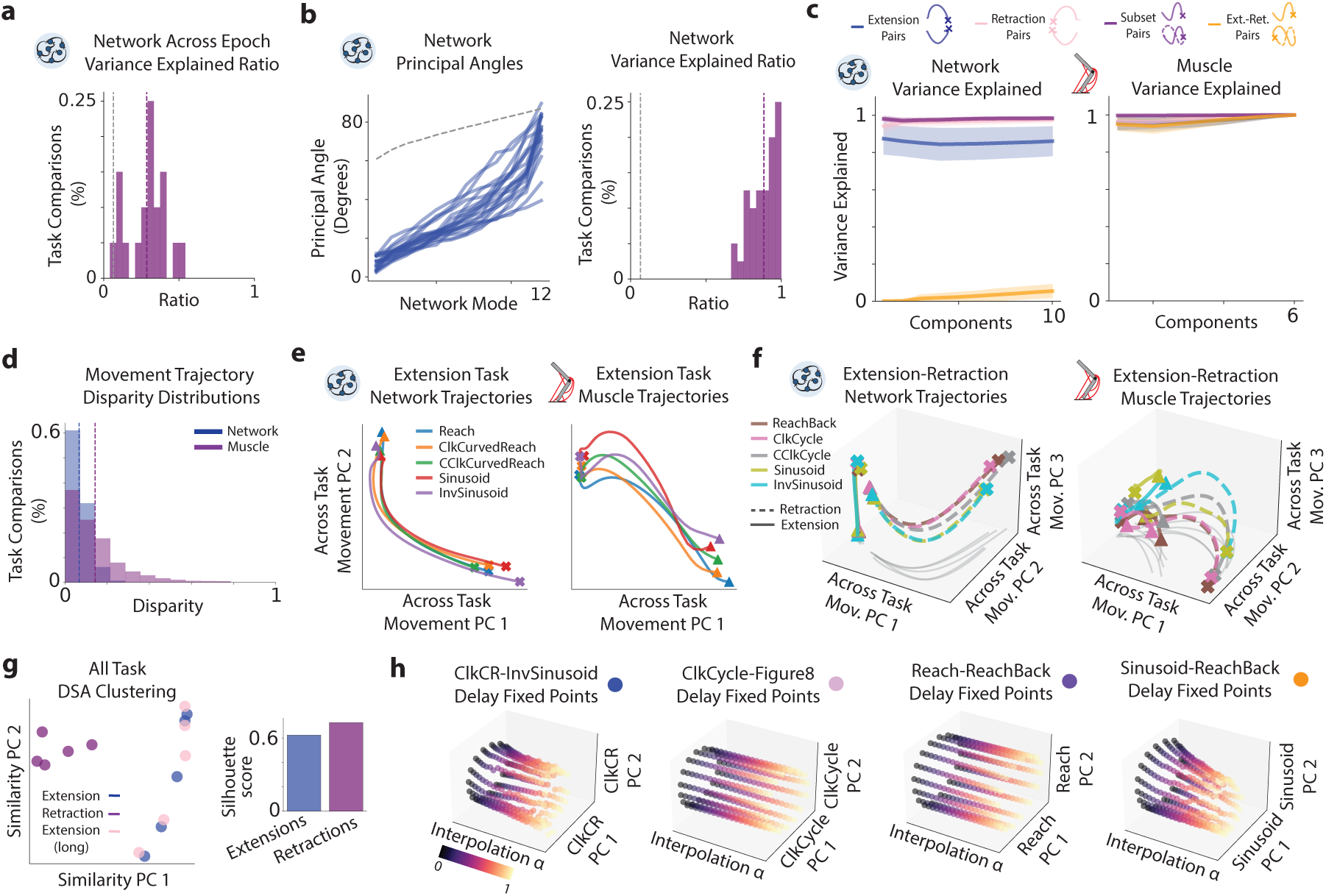
Shared low dimensional representation between similar movements across tasks. (a) Variance explained ratios across epochs within the same task shows orthogonal decomposition of preparatory and movement subspaces. (b) (left) Principal angles for the top 12 PCs (explaining over 95% of the variance) across similar tasks (extensions with extensions and extension-retraction with extension-retraction). Alignment across 12 components for all comparisons are below the random orthonormal baseline. (right) Ratio of variance explained for network activity projected onto the top 12 PCs of a different task compared to its own. (c) (left) Variance explained ratios as the number of PCs increase for different sub-movements in the network space, examples of which are shown at the top. All sub-movement pairs utilize shared subspaces except for extension and retraction pairs, which are implemented in orthogonal spaces. (right) Same as (left) but in the muscle activation space. Despite orthogonality in the network space for extension and retraction sub-movements, this distinction is not present in the muscle space. (d) Network and muscle disparity across sub-movements. Both systems show strongly aligned trajectories, with the network having stronger alignment than the muscles. (e) (left) Example network trajectories plotted onto the top two PCs found during movement. (right) same as (left) for muscle trajectories. (f) (left) Network trajectories plotted onto the first three PCs found during movement for extension-retraction tasks. (right) same as (left) for muscle trajectories. (g) (left) DSA on network trajectories during movement shows clustering of extension and retraction dynamics. (right) quantification of clustering through silhouette scoring. (h) Fixed point structures plotted onto the activity of task *A* as the network input is interpolated towards another task input. Fixed point structure largely remains the same throughout interpolation, showing shared use of fixed point structures across tasks.

Given that the network isolates computations in this way, one simple solution to the multitasking problem would be to isolate the tasks themselves: ten movements, each generated in its own subspace, with no interference between them. Compositionality may require the opposite. If the network is to build movements from reusable primitives, then tasks that share computations, for instance the five extension movements, should also share neural modes. To distinguish these possibilities, we compare the movement-epoch subspaces of different tasks using two complementary measures. Principal angles compare the dominant modes of two tasks: small angles mean the tasks occupy the same subspace and reuse the same modes, while angles near ninety degrees mean the subspaces are orthogonal. The variance-explained ratio, defined above, tests the same alignment in terms of the activity itself, measuring how well one task’s principal components account for the other task’s variance. We compare every measurement against a random-orthogonal baseline that accounts for the differing dimensionality of the network and the muscles, and we compute both measures between pairs of similar tasks (extensions with extensions, and extension-retractions with extension-retractions) using the top twelve PCs, which explain over 95% of movement-epoch variance.

We find that the network does not isolate tasks. Across task pairs, the principal angles are low relative to the random-orthogonal baseline (Fig. 3b, left), and the variance-explained ratios sit near one for task comparisons but near zero for the baseline (Fig. 3b, right). Similar movements are therefore generated within a shared set of neural modes, in line with reports of shared manifolds across tasks in primate motor cortex [20]. Note that neither metric requires the same numbered PC to align across tasks; they test whether the subspaces overlap, not whether PC 1 of one task matches PC 1 of another. A separate analysis of task variance shows that distinct unit populations are nonetheless recruited across tasks (Supplemental Fig. 1), indicating functional sub-structure within this shared manifold. Additionally, networks with varying hyperparameters such as activation functions and unit numbers consistently utilize shared low-dimensional manifolds (Fig. 3).

The globally shared manifold hides sharper structure at the level of sub-movements. Splitting extension-retraction tasks into their extension and retraction sub-movements, we plot the variance-explained ratio as a function of the number of components for four pair types: subsets (same kinematics, different rule input), extension pairs, retraction pairs, and extension-retraction pairs (Fig. 3c, left; see Methods). Subset, extension, and retraction pairs are all well aligned, with ratios near one, so the same subspaces are reused within each category. Extension-retraction pairs are the exception: their ratios stay near zero, meaning extensions and retractions unfold in nearly orthogonal subspaces. This contrast is absent in the muscle space, where extension-retraction pair ratios sit near one (Fig. 3c, right), indicating a principled separation of computations in the network that is not present in the arm. We attribute this to the much lower dimensionality of the muscle space, six muscles versus 256 units, which leaves less room to segregate computations into orthogonal subspaces. Keeping extensions and retractions in separate subspaces may be what allows them to be recombined freely, a possibility we return to when testing sequencing in the next section. Trajectory shapes, in contrast, are well aligned across all movement pairs in both spaces (Procrustes disparity close to zero), with stronger alignment in the network than in the muscles (Fig. 3d).

Next, we visualize the trajectories themselves. Across extension tasks, network and muscle trajectories overlap and share similar shapes when projected onto the top PCs (Fig. 3e). Extension-retraction tasks show a starker contrast between the two spaces (Fig. 3f): network trajectories move along PC 3 during extensions while retractions primarily occupy PCs 1 and 2, whereas muscle retractions traverse the same subspace as extensions in the opposite direction with a slight translation. These trajectories illustrate the result of Fig. 3c, namely distinct subspaces for extensions and retractions in the network and overlapping subspaces in the muscles.

Sharing a subspace shows that two movements use the same neural modes, but not that they use the same dynamics, since similar trajectories can be produced by different mechanisms [43]. We test for shared dynamics in two ways. First, during the movement epoch, we apply dynamical similarity analysis (DSA), which compares the vector fields of two systems directly, across all pairs of extensions and retractions [43]. The dynamical distances cluster by sub-movement type, with extensions and retractions forming separate groups (Fig. 3g, left), quantified by silhouette scores of around 0.6 for each cluster (Fig. 3g, right). Movements that share a subspace therefore also share their dynamics. Second, during the delay epoch, we ask whether the fixed point structure that prepares one task is reused to prepare another. The rule input is a static vector that selects the task, so we linearly interpolate it from task *A*’s value to task *B*’s value and recompute the network’s fixed points at each step (see Methods). If the two tasks rely on one dynamical landscape, the ring of preparatory fixed points should deform smoothly as the input changes; if they rely on separate solutions, the ring should collapse and be rebuilt somewhere else in state space. For four example pairs spanning the sub-movement categories of Fig. 3c, the ring-like fixed point structure is preserved throughout the interpolation (Fig. 3h. Together, the reuse of subspaces and dynamical features across tasks indicates a simplified solution to the multitasking problem, in which orthogonal subspaces separate distinct computations and shared manifolds are reused wherever the computation is similar.

### 2.4 Movements can be composed algebraically and sequentially through rule inputs

Shared subspaces and dynamics show that the network reuses the same neural modes across tasks, but not how the dynamics within these modes are combined to produce a particular movement. Nor do they show whether the extension and retraction motifs, which occupy separate subspaces, can be recombined into movements the network never explicitly learned. We address these questions through the rule input, the only signal that distinguishes one task from another. The rule input is a one-hot vector that stays constant throughout a trial and enters the network only as a constant input term, equivalent to a task-dependent bias. It carries no time-varying information and leaves the recurrent weights, which define the network’s dynamics, unchanged. Its sole effect is to bias the network into a different region of state space, and therefore to select among the fixed points and trajectories that the recurrent circuit already supports. If different tasks are produced by recombining the same primitives, then this shift should be enough: placing the network activity at the right location on a shared landscape, rather than rewiring it, should select the movement. First, we ask whether tasks are arranged compositionally in the delay subspace, and how the rule input positions the network within it. We then explicitly test if rule inputs can additively combine network primitives from a subset of movements to perform a movement outside the chosen subset. Second, we test sequencing by switching the rule input mid-movement, chaining the extension motif of one task with the retraction motif of another, a pairing the network never encountered during training. Together, these tests ask whether the network’s computations are composable, that is, whether novel trajectories can be produced from learned parts by nothing more than a change of position in state space.

To look for explicit compositional representations, we apply targeted dimensionality reduction (TDR; see Methods) to the condition-averaged activity during the delay epoch to discover axes distinguishing the rotational and reach frequency components of the five extension tasks (Fig. 4a). The tasks fall into a structured grid. The rotation axis well separates clockwise and counter-clockwise movements, with straight reaches situated between both. The frequency axis distinguishes reaches, curved reaches, and sinusoids in increasing order. This demonstrates a compositional strategy for initializing movement. Interestingly, we find that varying network size with different activation functions does not always lead to a structured grid; however, Softplus networks consistently contain this representation (Fig. 3).

**Fig. 4.**
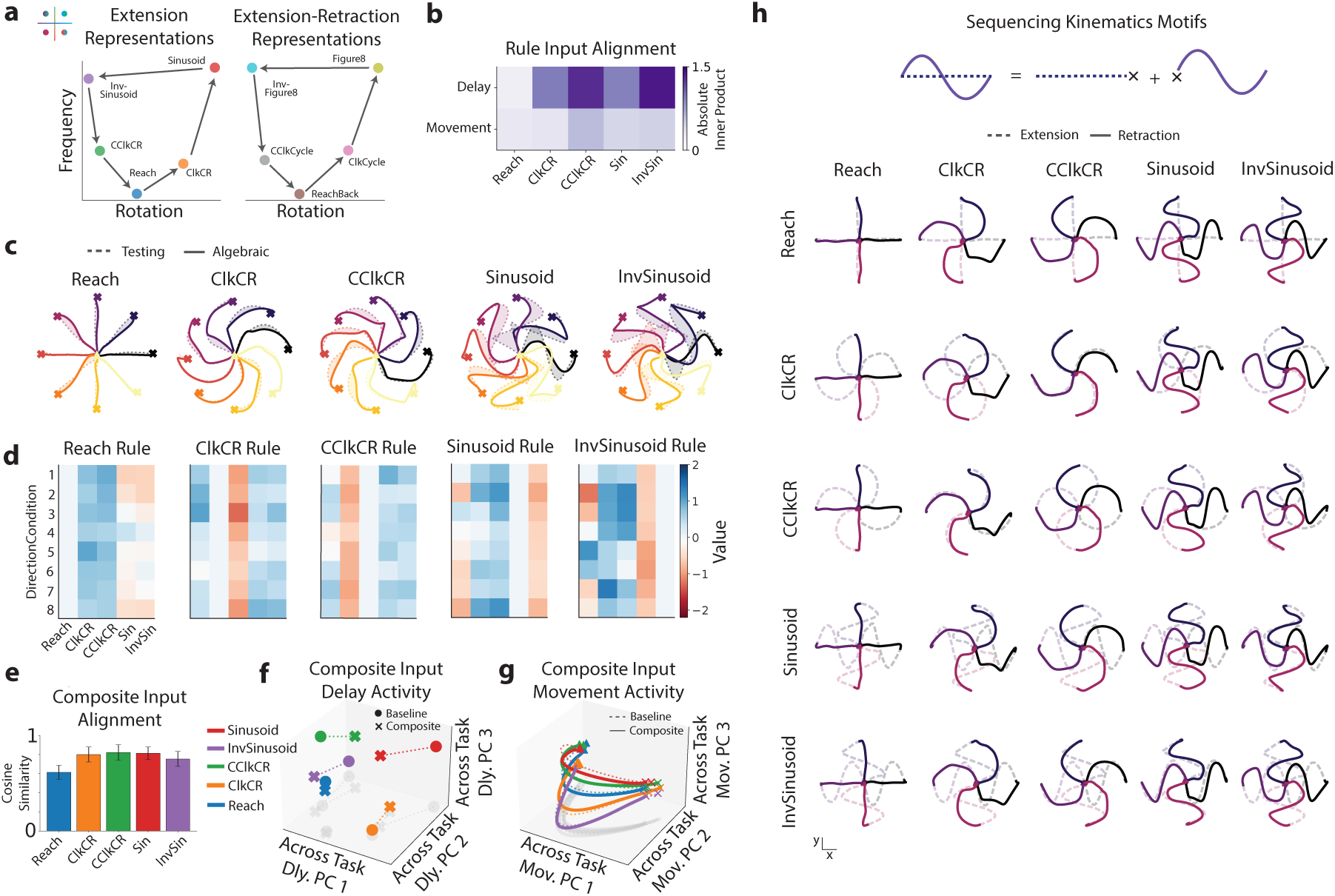
Compositional representations. (a) Algebraic representation of tasks during the delay epoch, found using TDR. Rotation and frequency axes are mostly orthogonal and offset tasks in a structured manner. (b) Rule input alignment with the delay and movement subspaces found by taking the rule input weights and computing the weighted average of the absolute inner product across PCs. The first five and the first 17 PCs are used for the delay and movement subspaces respectively. (c) Network kinematics on novel rule input found through an algebraic composition of other tasks. Each task is able to perform similar to its baseline performance despite not using the original rule input. (d) Exact value of the composite rule inputs used in (c) across the eight direction conditions. (e) Cosine similarity between composite input vectors and original rule input vectors for each task. Input vectors are found by projecting the rule inputs (composite or original) into network space. Both the composite and standard rule vectors are further projected onto the first 18 PCs found across tasks, conditions, and epochs, to ensure only task relevant modes are considered. (f) Initial condition during delay using original rule input and composite rule input, plotted on the PCs captured using the original rule input. (g) Same as (f) but during movement. (h) Sequencing kinematic trajectories by switching the rule input after an extension task demonstrates the network’s ability to generate new movements by stitching together distinct extensions and retractions. Rows denote the extension task and columns the retraction task; dashed lines show the extension segment and solid lines the retraction segment of each hand trajectory. Each combination of extension and retraction performs well across multiple directions despite never explicitly training on these movements.

As previously discussed, each task is distinguished by a static rule input, which acts to shift the dynamical landscape of the network to appropriately perform a desired task. There are two potential strategies to achieve this: the rule input could primarily shift the delay state to properly initialize movement, relying on generalized movement-related dynamics, or it could also directly shift movement dynamics to accomplish distinct tasks. We test between these possibilities with a rule vector alignment metric across the delay and movement subspaces (see Methods). We find that the rule input is strongly aligned with the delay subspace across most tasks, and less so with the movement subspace (Fig. 4b). This suggests that the rule input primarily shifts the state in the delay subspace to prepare movements in a principled manner.

With insight as to how the network prepares movement and how the rule input helps accomplish this, we ask if rule inputs can be combined to obtain the correct primitives for distinct movements. We test this using an optimization procedure that finds combinations of existing rule inputs to perform each task. Crucially, no network weights change: the recurrent, input, and output weights all stay frozen, and we search for a combination of the other tasks’ rule inputs that drives the network to perform a held-out task. The network finds such a combination for every task, and output kinematics closely match the target task (Fig. 4c). The explicit values for the rule inputs are shown in Figure 4(d); for each task its own rule input value is set to zero, while the values for other tasks are specific combinations of positive or negative values for appropriate offsetting. Because the circuit is unchanged and only the input is recombined, this shows the internal computations are composable, not just that the geometry has a qualitative compositional structure.

Composite rule vectors may achieve task performance by closely aligning with original rule vectors, or through more complex mechanisms and individualized solutions. To test alignment, we take the cosine similarity between the composite rule vector, found by taking the optimized composite rule input and multiplying it with the corresponding input weights, and the standard rule vector. We find that composite rule vectors are strongly aligned with original rule vectors across tasks and conditions, with cosine similarities around 0.8 (Fig. 4e). Upon closer inspection, we see that the initial condition of the network before movement (last timestep of delay epoch) is close to that of the original rule input, with some misalignment along PC 2 (Fig. 4f). Additionally, movement trajectories with composite inputs closely overlap with original movement trajectories (Fig. 4g). This shows that the optimal composite rule vector for a given task is one that is close to the original rule vector, and the network is able to find this with additive combinations of other task inputs.

The second form of compositionality we consider is sequencing. During training, each extension is paired only with its own retraction, for example a reach followed by a reach back, so the network never sees an extension from one task followed by a retraction from another. We ask whether the extension and retraction motifs are nonetheless stored as separate units that can be recombined. To test this, we activate an extension task’s rule input and then, mid-movement, switch to the rule input of a different extension-retraction task, so that a retraction the network has never performed after this extension is requested by a change in the static input alone (Fig. 4h). Most task pairs execute the novel extension-then-retraction combination accurately. Since the network was never trained on these pairings, its ability to produce them implies that the extension and retraction motifs are reusable units rather than components of a single, task-specific trajectory. It is this modularity, rather than the ordering of two movements in time, that we take to be compositional, consistent with how sequential primitives are defined in the motor-control literature [13, 37, 44].

### 2.5 Embodied, feedback-driven training constrains the network towards compositional solutions

The compositional structure described so far emerged in a network that controls a body. Unlike cognitive multitasking models, whose outputs directly represent abstract decision variables, our network produces high-dimensional muscle commands that drive an arm, and it receives time-varying feedback from that arm. Whether this setting helps or hinders compositionality is not obvious. Task relationships that are evident in hand kinematics need not be evident in muscle activity, because the arm is redundant: many distinct muscle commands produce the same hand trajectory. A network could therefore learn a separate muscle-level solution for each task, with no shared structure across tasks. We ask two questions. First, does this redundancy permit non-compositional solutions, and if so, why did the baseline network not find one? Second, which components of the embodied setting constrain the network towards reusable primitives? To answer them we train three control models (Fig. 5a,b). The *Niche Solutions* model is trained to reproduce the muscle activations of ten separate networks, each trained on a single task. The *No Feedback* model is trained on the standard end-effector loss but without proprioceptive or visual feedback. The *No Effector* model receives the same inputs as the baseline network and is trained to reproduce the baseline network’s muscle activations directly, without the arm in the training loop. Each of the three models performs all tasks well (Fig. 5c-e). For each, we test algebraic composition through the rule input as in Section 2.4, and use the analyses introduced above to identify where composition degrades and why.

**Fig. 5.**
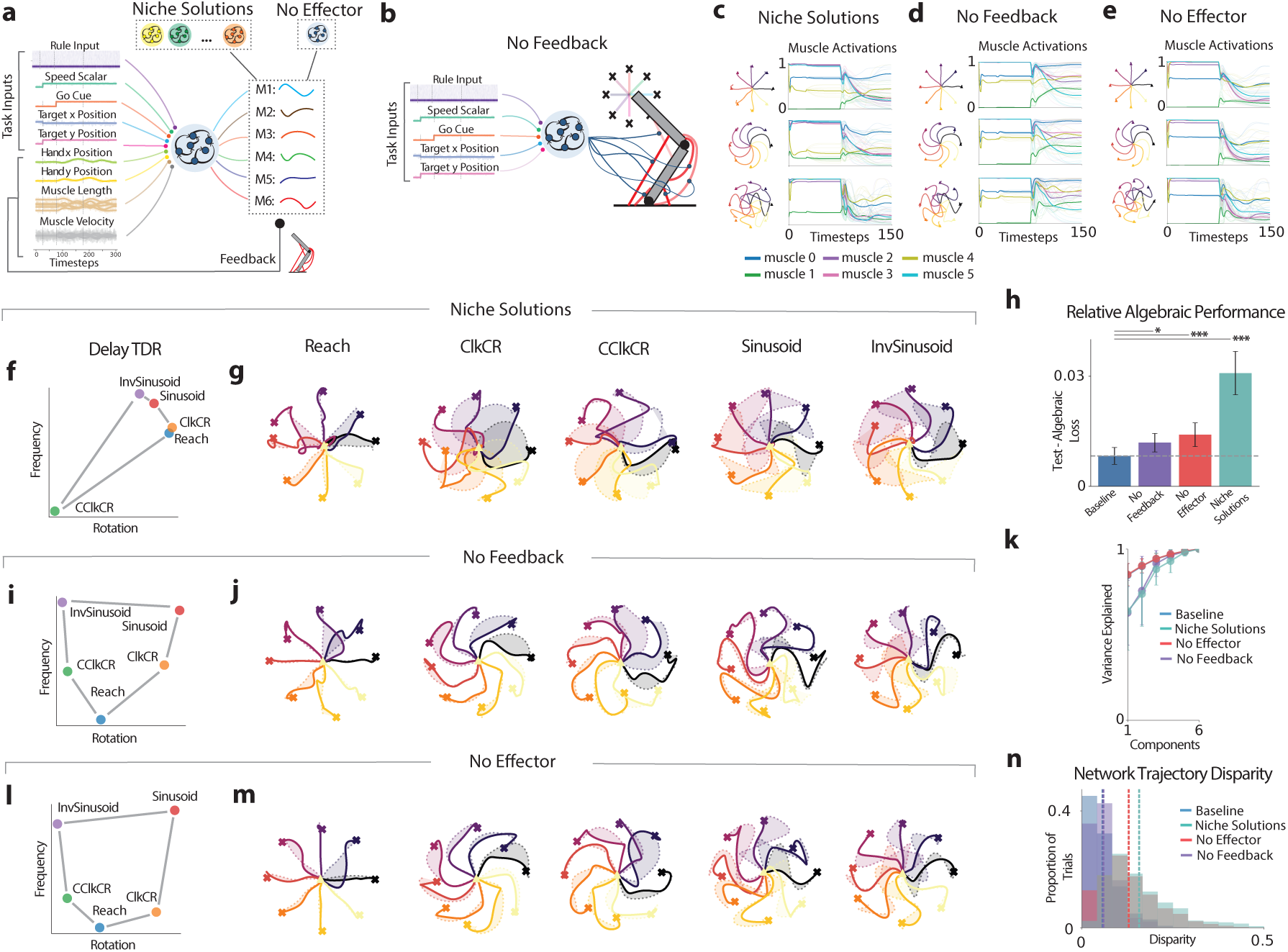
Ablations of feedback, embodiment, and embodied multitasking. (a) Schematic of Niche Solutions and No Effector model, which train to directly regress muscle activations while feedback is provided. The Niche Solutions model is trained using muscle activations found by ten distinct networks trained on each task individually, while the No Effector model is trained using muscle activations found by the baseline network. (b) Schematic of the No Feedback model, which is trained with the standard end effector loss but without the use of proprioceptive and visual feedback. (c) Example kinematics and muscle solutions for the Niche Solutions model. (d-e) Same as (c) for the No Feedback and No Effector models respectively. All models can perform individual tasks well. (f) TDR on network delay activity shows lack of structured geometry across tasks for the Niche Solutions model. (g) Kinematics with algebraic rule input for the Niche Solutions model, dotted lines denote the kinematics with the original rule input for that model. (h) Relative algebraic performance for each model, which is the difference in MSE found between the kinematics performed using the standard rule input for each task subtracted and the algebraic rule input. Significance found using t-test between baseline and other models. (i-j) Same as (f-g) for the No Feedback model. (k) Variance explained across components in the muscle activation space across models. The No Feedback and Niche Solutions networks are the highest dimensional. (l-m) same as (i-j) for the No Effector model. (n) Procrustes on network trajectories across tasks for different models. The No Effector and Niche Solutions models have the largest disparity in the network space.

We first establish that redundancy alone permits non-compositional solutions. The Niche Solutions model performs every task in kinematic space, but each task is produced by a muscle pattern found independently of the others. This network shows no compositional structure: the delay-epoch geometry found by TDR is unstructured (Fig. 5f), and composite rule inputs fail to produce the target movements (Fig. 5g). To compare models, we compute the relative algebraic performance as the difference in kinematic loss between the standard and composite rule inputs (Fig. 5h). The Niche Solutions model shows the largest degradation; the other two models fall closer to the baseline but remain significantly worse. The Niche Solutions model also uses the rule input differently from the baseline network: its rule input is aligned with the movement subspace to a far greater degree than in the baseline network, rather than being confined to the delay subspace, and its composite rule vectors are poorly aligned with the original ones (Supplemental Fig. 4). Redundancy in the mapping from network output to kinematics can therefore produce a solution with no compositional structure, which implies that multitask training through the arm steered the baseline network away from such solutions.

We next ask which components of embodied training provide this constraint. The remaining two models each remove a single component from the baseline: the No Feedback model removes the sensory feedback from the arm, and the No Effector model replaces the loss on end-effector kinematics with a loss on muscle activations. Without feedback, the network still develops a structured task geometry during the delay epoch (Fig. 5i) and can compose tasks from rule inputs (Fig. 5j), but it does so less accurately than the baseline, and its composite rule vectors are less aligned with the standard ones (Supplemental Fig. 4). One difference in its solution is dimensionality: the No Feedback model uses higher-dimensional muscle activation patterns than the baseline (Fig. 5k). A higher-dimensional mapping from network to muscles leaves more room for task-specific solutions, which may be what limits composition in this model.

Trained on muscle activations rather than kinematics, the No Effector model has the same inputs and outputs as the baseline but is no longer required to control the arm. Like the No Feedback model, it develops a structured delay-epoch geometry (Fig. 5l) but composes tasks less accurately than the baseline (Fig. 5m), with composite rule vectors whose alignment to the standard ones varies strongly across tasks (Supplemental Fig. 4). Here the difference lies in the network trajectories: Procrustes disparities across tasks are considerably larger than in the baseline (Fig. 5n), so the No Effector model does not reuse trajectory shapes across tasks even though it reproduces the baseline’s outputs. Training the network to control the arm, rather than to reproduce a set of muscle patterns, therefore constrains its solutions towards shared trajectories and stronger compositional structure. Together, the three models show that the baseline network’s compositionality depends on both sensory feedback and control through the arm: removing either degrades it, and replacing the shared multitask solution with independent, task-specific muscle solutions eliminates it.

### 2.6 Transfer learning for novel movements

A compositional system should be able to acquire a new but related movement quickly, by recombining the primitives it already has rather than learning a new solution from scratch. We test this with transfer learning. We pre-train the network on eight of the ten tasks, then freeze all recurrent, input, and output weights and allow only the input weights of the two held-out rule inputs to be learned. As in the previous sections, the rule input can only bias the network into a different region of state space, so if the held-out movements can be learned in this way they must be assembled from motifs that the network already possesses. We then vary which tasks are included in pre-training to ask which of those motifs the held-out movements depend on.

We hold out the counter-clockwise curved reach (CClkCR) and the counter-clockwise cycle (CClkCycle). With only their two rule-input weight vectors free to change, the network learns both tasks nearly perfectly (Fig. 6b,c).

**Fig. 6.**
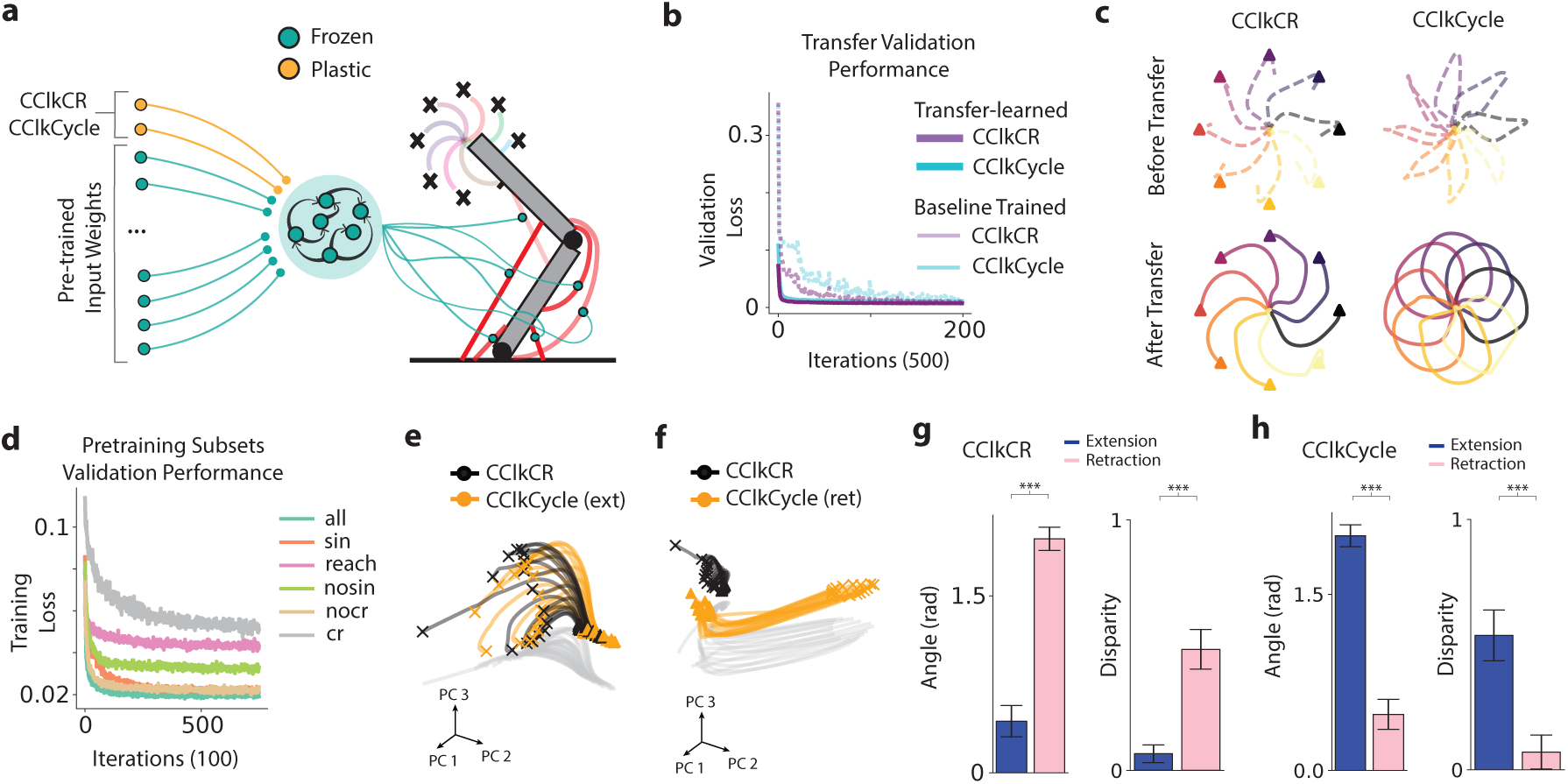
Transfer of a pretrained model towards novel tasks. (a) Task setup. The model is pretrained on all tasks except for CClkCurvedReach (CClkCR) and CClkCycle, and all but the input weights for the rule inputs for these two tasks are frozen in the transfer setting. (b) Validation loss during transfer learning on the held-out tasks. Dashed lines denote baseline model performance on same tasks. (c) Example kinematics for both tasks before (top) and after (bottom) transfer learning. (d) Networks pretrained on fewer task subsets were not able to perform the held-out transfer learning tasks as well as those containing all possible task subsets. Training MSE loss shown during transfer learning of each pre-trained model. (e) Network trajectories for CClkCR and the extension portion of the CClkCycle task projected onto the top three CClkCR PCs. (f) Same as (e) but for the retraction portion of the CClkCycle task. (g) Angular distance and disparity metrics from the CClkCR task to all other extension and retraction tasks. CClkCR trajectories are better aligned with the extension tasks as opposed to retraction tasks in both metrics (p *<* 0.001, angular distance; p *<* 0.001, disparity; Mann-Whitney U test). Error bars denote standard deviation across pairs. (h) Same as (g) but for the retraction portion of the CClkCycle task. As expected, this task is closely aligned with other retraction tasks (p *<* 0.001, angular distance; Mann-Whitney U test).

We next test how transfer depends on the pre-training set (Fig. 6d). We evaluate six pre-training subsets, none of which includes the held-out tasks: all eight remaining tasks (all); only the sinusoid and figure-eight tasks (sin); only Reach and ReachBack (reach); all tasks except the sinusoids and figure eights (nosin); all tasks except ClkCR and ClkCycle (nocr); and only ClkCR and ClkCycle (cr). Restricting the pre-training set degrades transfer, and the full set transfers best. Pre-training on Reach and ReachBack alone, for example, does not enable curved reaches, so the held-out tasks require computations that must be assembled from several learned primitives rather than inherited from a single related task. Whether richer natural movements could be built from such primitives is a hypothesis we leave for future work.

The transfer solutions reuse the pre-trained subspace structure. During movement, CClkCR trajectories overlap with the extension phase of CClkCycle (Fig. 6e) but not with its retraction phase (Fig. 6f), echoing the extension-retraction orthogonality found in Section 2.3. Quantifying alignment with angular distance and Procrustes disparity, CClkCR aligns with the other extensions (Fig. 6g), while the CClkCycle retraction aligns with the other retractions (Fig. 6h). The network thus solves the novel tasks by reusing previously learned extension and retraction subspaces, a hallmark of compositional representation.

## 3 Discussion

Flexible, skilled movement requires the ability to learn and deploy a wide repertoire of actions without learning each one from scratch. To understand how a single network can reconfigure its computations across many movements, we designed a model and task suite that allow systematic analysis of shared computations across contexts. RNNs have become a standard tool for hypothesis testing in motor control and have yielded biologically informative predictions [25, 45]; more recently, they have been extended to include biomechanical feedback [30, 31, 35], an integral component of the motor system [46]. Our tasks mirror the center-out paradigms used in primate experiments, and the sub-movements are simple and well defined, so that network representations and shared computations can be compared directly with experimental findings.

Our model sits within the framework of optimal feedback control (OFC), in which movement is generated by a feedback policy that maps the current sensory state to motor commands so as to minimize a task cost [46, 47]. The trained network is such a policy, learned rather than derived: it receives delayed visual and proprioceptive feedback and is optimized on end-effector error. Recurrent networks trained in this way have been shown to reproduce signatures of OFC and of motor cortical activity [48, 49]. Because feedback is delayed and the loss is defined on the hand, the network must anticipate the sensory consequences of its commands; the predictive computation that internal-model theories assign to a forward model [50] is therefore present, but embedded in the recurrent dynamics rather than implemented as a separate module. Our analyses do not attempt to isolate forward and inverse components, and we make no claim about that decomposition. What the results do address is how a single feedback controller organizes many movements. Modular theories such as MOSAIC propose multiple paired internal models selected by context [50, 51]. We find a different organization: one shared dynamical landscape, within which a static task input selects a movement by relocating the network state, with modularity appearing only at the level of extensions and retractions, which occupy orthogonal subspaces. OFC also predicts that task-irrelevant dimensions of a redundant plant are left uncontrolled [46]. Our Niche Solutions model shows that this redundancy could support a separate muscle-level solution for every task, yet the network trained through the arm with feedback converged on shared, low-dimensional muscle patterns instead. OFC specifies the optimum but not the path learning takes to reach it; our ablations suggest that learning a feedback policy through the plant constrains that path towards reusable solutions. Finally, the transfer results show that a new movement can be acquired by re-parameterizing the input to a fixed policy, the counterpart in a learned controller of changing the goal of an optimal one. The network is best read as a lumped model of sensorimotor circuitry rather than M1 alone, and it does not address higher cognitive processes such as mental simulation, though the delay epoch captures a basic preparatory computation.

In recent years, there has been a shift from studying the tuning of individual units to examining population-level dynamics during the control of movement [52]. This shift has revealed low-dimensional manifolds, and with them a view of movement generation as the unfolding of population dynamics along a small number of neural modes that carry the task-relevant computations [25]. In our model, distinct computations, whether across epochs or between extensions and retractions, occupy orthogonal subspaces, while similar movements share the same subspaces and the same dynamics. The shared structure is in line with experimental findings of task-invariant neural modes in motor cortex [20], and suggests that commonalities between movements are represented by a common set of modes that serve as building blocks for movement generation. The network thus either reuses or orthogonalizes subspaces depending on the computational similarity of the movements involved.

Shared subspaces and dynamics provide the substrate for compositional representations. The network arranges tasks algebraically in the delay subspace, separating rotational direction from kinematic complexity. We use a broad notion of compositionality, consistent with the computational neuroscience literature: a representation is compositional if it is built by combining multiple reusable components, whether those components are discrete or vary continuously [8, 9, 53]. Importantly, the evidence for compositionality is functional and not only geometric. With all weights frozen, additive combinations of other tasks’ rule inputs drive the network to perform a held-out task, and the resulting composite inputs closely approximate the original rule input for that task. Because the rule input can only bias the network to a different location in state space, this shows that the dynamics required for each movement are already present in the shared circuit and are selected, not constructed, by the task input.

Sequencing provides a second, independent line of evidence for compositionality. The network produces extension-retraction combinations it was never trained on, which means it stores extension and retraction motifs as separate, reusable units rather than as one bespoke representation per compound task. The compositional content lies in this modular reuse, not in the ordering of two movements in time, in keeping with definitions of motor primitives in the literature [37, 44]. The orthogonal subspaces and distinct dynamics we find for extensions and retractions offer a mechanistic basis for this modularity: because the two motifs are implemented in separate dynamical regimes, switching the rule input mid-movement can recruit a new retraction without disrupting the extension that preceded it.

The role of the body in shaping neural representations is an increasingly central question in motor neuroscience. The arm is redundant, since many muscle trajectories can produce the same kinematic trajectory, so a multitasking network could in principle adopt an independent muscle-level solution for each task. Our Niche Solutions model shows that such solutions exist and that they carry no compositional structure. The baseline network did not converge on one, which suggests that embodied multitask training constrains the solution space, and the two remaining ablations identify which components of embodied training are necessary for compositional structure to emerge. Removing sensory feedback leads to higher-dimensional muscle activity and weaker composition, suggesting that feedback constrains the network towards low-dimensional, shared output patterns. Removing the arm from the training loop, while keeping inputs and outputs identical, leads to more varied network trajectories across tasks and again weaker composition, suggesting that the requirement to control the hand, rather than merely to reproduce muscle patterns, constrains the network towards shared trajectories. Embodied training therefore does not merely permit compositional representations; it steers the network towards them in a setting where non-compositional solutions are readily available. One significant advantage of compositional representations is the ability to quickly adapt in novel settings. Multitasking models of cognitive tasks learn new tasks faster after pre-training on related ones [54] and can acquire new tasks using previously learned motifs alone [9]. Our network shows the analogous capability for movement: with all weights frozen, a new rule input suffices to produce a held-out movement from existing extension and retraction subspaces, provided the relevant primitives were present during pre-training. This parallels the finding that primates rapidly learn within-manifold perturbations of a center-out task [55].

One limitation of this model is the relatively low number of muscles in the arm, along with its planar two-dimensional structure. It is possible that with greater redundancy, i.e., more muscles and a higher-dimensional task space, there may be a more complex relationship between network and muscle modes. Additionally, it may be hypothesized that the simple low dimensional manifolds typically found during lab experiments, as well as in our setting, may reflect the simplicity of the tasks themselves. The addition of full body musculoskeletal models capable of performing a wide array of naturalistic behaviors may enhance our understanding of the underlying motor manifold, which may be non-linear and higher dimensional in nature. Hierarchical structures in the motor system likely also play a vital role in understanding the compositional nature underlying motor control. Framing the motor cortex as a high-level controller may abstract the representations found in this study to instead depend on the combination and interaction between low-level controllers. Despite the simplicity of our current setup, our framework was able to clearly identify both shared and isolated computations in addition to compositional structure used by networks to achieve flexible control of movement, and can be adapted in the future to include more complex musculoskeletal models and a wider variety of tasks reflecting less stereotyped paradigms.

## 4 Methods

### 4.1 Network and arm structure

We train standard RNNs in feedback with a musculoskeletal model to perform ten unique movements designed to elucidate a compositional structure in the network activity (Fig. 1a,b). The form of the network is described below:

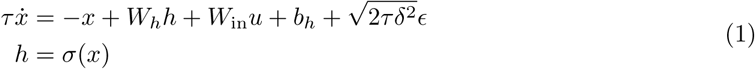

We define every symbol at first use and state whether it is a scalar, vector, or matrix. Here the vector *x* is the pre-activation state (the membrane-potential-like variable before the nonlinearity), the vector *h* = *σ*(*x*) is the firing rate, and *σ* is the network activation applied element-wise. The matrices *W_h_* and *W*_in_ are the recurrent and input weights, the vector *b_h_* is the bias, the vector *u* is the input, and the vector *ɛ* ∼ *N*(0, *I*) is Gaussian noise. The scalar *δ* = 0.1 sets the noise magnitude and *τ* = 20 is the network time constant. We add noise to the pre-activation to model biological variability and to improve the robustness of the trained solution, a standard practice in RNN models of neural circuits [45, 56]. The factor 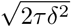 comes from the continuous-time (Ornstein-Uhlenbeck) formulation: it keeps the stationary noise variance fixed as the discretization step Δ*t* changes, so *δ* is a single interpretable noise scale rather than a step-dependent quantity.

We discretize 1 in order to train the network using conventional deep learning methods; the form of the discretized network is given as

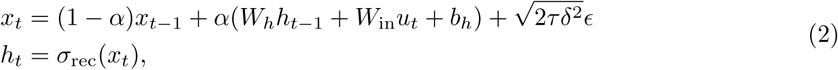

The term *α* = Δ*t/τ*, where Δ*t* = 20. We utilize a readout layer that transforms network outputs to muscle stimulations in the range (0, 1)

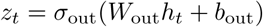

In this case *σ*_out_ is the sigmoid function 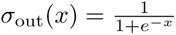. The output layer formulation is consistent across all hyperparameter settings due to the requirements of the arm.

For the musculoskeletal model we use Motornet [29], an open-source framework for designing musculoskeletal models with differentiable muscles in Python. This allowed us to directly backpropagate through the muscles and train the network using supervised learning. The model itself contains six Hill-type muscles and two joints, capturing horizontal movements at the shoulder level. Each muscle represents a lumped representation of synergistic muscle groups in a primate arm, with a total of 19 muscles considered [39].

### 4.2 Task formulation and training

We use “multitasking” in the sense established in the cognitive and machine-learning literature: a single network trained to perform several distinct tasks [8, 9]. It does not refer to performing two tasks simultaneously or to dual-task interference. We define the term at its first use in the Introduction and retain it here for consistency.

For each task the arm starts in the same initial joint state, representing a center-out setup paradigm (Fig. 1a), then is instructed to move in one of sixteen directions at one of five different speeds. For validation, we save the model that has the best performance on held out, interpolated conditions including 32 directions and 10 speeds. The RNN receives proprioceptive feedback in the form of muscle lengths and velocities, and visual feedback of the end effector position (Fig. 1a). Other task-related inputs include a speed scalar, coordinates of the target position, go cue, and one-hot rule input. A trial is one execution of a single task, in which the network moves the arm in one direction at one speed. Each trial has four sequential epochs (Fig. 1c): (1) baseline, where the arm and network settle to rest; (2) delay, where the network receives the upcoming movement’s direction, speed and task identity (the rule input) but must not move, analogous to the preparatory period in primate motor experiments; (3) movement, where the arm executes the instructed trajectory; and (4) hold, where the arm maintains its final position. A condition refers to a particular direction-and-speed setting of a task. These definitions also appear in the main text (Results, Section 2.1) before any result that uses them. For each task, the length of the baseline *T*_baseline_ and hold *T*_hold_ epochs are *T*_baseline_ = 25, *T*_hold_ = 25. For the delay epoch, the length is randomly selected according to *T*_delay_ ∼ {50, 75, 100}. The length of the movement epoch during training is determined by the chosen speed condition, and ranges from *T*_move_ ∼ {50, 75, 100, 125, 150} for extension only tasks and *T*_move_ ∼ {100, 150, 200, 250, 300} for tasks containing extension and retraction.

The kinematics of the tasks themselves are designed to contain compositional elements (Fig. 1b). The task suite is as follows: reach (Reach), clockwise and counter clockwise curved reach (ClkCurvedReach, CClkCurvedReach), sinusoid (Sinusoid) and inverse sinusoid (InvSinusoid), reach outward and backward (ReachBack), clockwise and counterclockwise cycles (ClkCycle, CClkCycle), and lastly figure eight and inverse figure eight (Figure8, InvFigure8). While each task in its entirety represents a unique movement pattern, there are various compositional elements. For example, the extension kinematics of the ClkCycle task are the same as the ClkCurvedReach task, certain tasks only involve extension, while some involve retraction, and some tasks are simply rotated versions of another (e.g. ClkCycle, CClkCycle).

The network is trained on five separate losses, given as

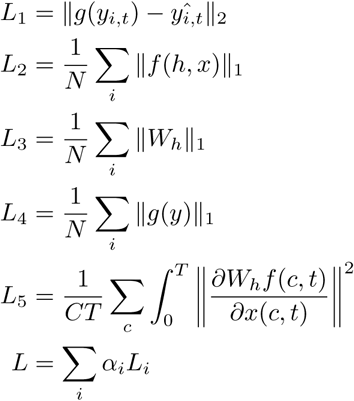

Here *g* maps muscle activity to hand kinematics (the musculoskeletal dynamics), *f* denotes the discretized RNN dynamics defined in Eq. 2, *W_h_* are the recurrent weights, and *x* is the pre-activation state (firing rate *h* = *σ*(*x*)). The first loss *L*_1_ is the mean squared error between hand position and the desired trajectory, and the network never receives this desired position as feedback. Losses *L*_2_, *L*_3_, and *L*_4_ are *ℓ*_1_ penalties on the network rates, recurrent weights, and muscle activations respectively. The loss *L*_5_ is a dynamical smoothness penalty [45]: intuitively, it penalizes rapid changes in the recurrent dynamics across time and so encourages smooth, slowly varying solutions, motivated by the smooth low-dimensional population dynamics seen in cortex during reaching [38]. Our total loss is the sum of each individual loss scaled by a weighting factor *α_i_*. For the base model shown in the paper we use *α*_1_ = 10^−3^, *α*_2_ = 10^−3^, *α*_3_ = 10^−3^, *α*_4_ = 10^−3^, *α*_5_ = 10^−3^. We train the model for 75, 000 iterations using a batch size of 32, and 256 units for the base model. The optimizer used is Adam [57], with a learning rate of 10^−3^.

For loss *L*_1_, we generate desired kinematics for each task. The methods for doing so are presented below:

- **Reach**: Generate a line from the initial hand position *z*_0_ ∈ ℝ^2^ to the target position *p̂* ∈ ℝ^2^: *t*^^^= *z*_0_(1−*α*) + *αp̂*, *α* ∈ [0, 1]. The target position is a random orientation on the unit circle *p* ∈ ℝ^2^, chosen from sixteen equally spaced bins, scaled and shifted by *β, δ* ∈ ℝ respectively to be centered at the hand position, *p̂* = *βp* + *δ*.
- **ClkCurvedReach**: Generate a scaled and shifted curved trajectory *t*^^^(*x_i_*) = *β*(cos(*x_i_*), sin(*x_i_*)) + *δ, β, δ* ∈ ℝ. In this case *x_i_* = *π*(1 − *α_i_*), 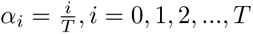 where *T* is the chosen number of timepoints.
- **CClkCurvedReach**: Same as ClkCurvedReach except *x_i_* = *π*(1−*α_i_*) + 2*πα_i_*.
- **Sinusoid**: Generate a scaled and shifted curved trajectory *t*^^^(*x_i_*) = *β*(*x_i_,* sin(*y_i_*)) + *δ, β, δ* ∈ ℝ, where 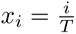, *y_i_* = 2*πα_i_,* 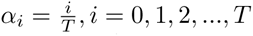.
- **InvSinusoid**: Same as Sinusoid except *t*^^^(*x_i_*) = *β*(*x_i_,* −sin(*y_i_*)) + *δ*
- **ReachBack**: We combine the trajectory used in the Reach task, *t*^^^_1_, with a separate trajectory from the target position back to the center: *t*^^^_2_ = *p̂*(1 *α*) + *αz*_0_. The total trajectory is defined as *t*^^^= [*t*^^^_1_, *t*^^^_2_].
- **ClkCycle**: We combine the trajectory used in the ClkCurvedReach task, *t*^^^_1_, with a separate trajectory *t*^^^_2_ in the same direction circling back to the initial condition using a distinct *x_i_*: *x_i_* = −*πα_i_,* 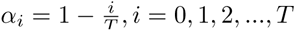. The total trajectory is defined as *t*^^^= [*t*^^^_1_, *t*^^^_2_].
- **CClkCycle**: Same method as ClkCycle, using the CClkCurvedReach trajectory as *t*^^^_1_, except the *x_i_* used for *t*^^^_2_ is given as *x_i_* = *πα_i_,* 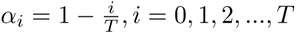.
- **Figure8**: We combine the trajectory used in the Sinusoid task, *t*^^^_1_, with a separate trajectory *t*^^^_2_ = *β*(*x_i_,* − sin(*y_i_*)) + *δ, β, δ* ∈ ℝ. Here, 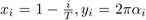, 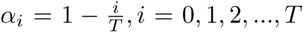. The total trajectory is defined as *t*^^^= [*t*^^^_1_, *t*^^^_2_].
- **InvFigure8**: Same method as Figure8, using the InvSinusoid trajectory as *t*^^^_1_.

### 4.3 Manifold Analysis

In order to compose previously learned motifs across tasks, the network may utilize a shared space in which the same network modes are reconfigured to perform similar movements in distinct settings. To test whether or not a shared low dimensional manifold exists across tasks, we utilize the methods presented in [20, 58] to compute the principal angles between network modes across tasks. Briefly, given *X_A_* and *X_B_*, the top *m* principal components (PCs) for tasks *A* and *B* respectively that explain above 95% of task variance, we compute the principal angles between these modes by computing *C* via 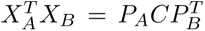 using singular value decomposition, where *X_A_* and *X_B_* are the *n* by *m* matrices containing the top *m* PCs for tasks *A* and *B*, and *P_A_* and *P_B_* are the *m* by *m* matrices defining the new manifold directions that minimize the principal angles. The essence of this method is to successively find modes that align with each other for both tasks. The matrix *C* contains the ranked cosines of the principal angles *C* = diag(cos(*θ*_1_), cos(*θ*_2_), *…,* cos(*θ_m_*)). In addition to principal angles, we also compute the ratio of variance explained between the PCs across and within tasks. The form of this ratio is simply var(*X_B_h_A_*)*/*var(*X_A_h_A_*). The baseline for both metrics is a random orthogonal *m* by *N* basis, built using a QR decomposition, where *m* is the number of basis vectors chosen for computing principal angles (in this case 12 for the network and three for the muscles). To define the number of similar modes across task and condition comparisons in Figure 3(b), we take the 0.1 percentile of 5000 comparisons of randomly generated bases to create a *P <* 0.001 threshold, below which the observed principal angles in the network and muscles are considered significant.

In Section 2.3, we group task pairs based on their similarities and dissimilarities then compute their angular distance to test their alignment. The angular distance is time-averaged as such

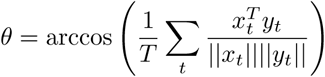

When grouping kinematic pairs, we separate extension-retraction tasks into two halves to compare both phases of the movement distinctly. The task pairs are as such:

- Subset: Tasks with different rule inputs but the same kinematics (e.g., Reach and the extension half of ReachBack).
- Extension: Any pair of extensions excluding subsets (e.g., ClkCR and Sinusoid)
- Retraction: Any pair of retractions, found from the second half of extension-retraction tasks (e.g. retraction phase of Figure8 and ClkCycle).
- Extension Retraction: Any extension paired with any retraction (e.g. Reach and retraction phase of Figure8).

### 4.4 Latent Analysis

In order to test the alignment of network and muscle latents, we utilize CCA and Procrustes analysis. CCA aims to find the modes such that the single-dimensional latent activity of two datasets *A* and *B* are maximally correlated. For CCA, we first compute the first ten PCs of task *A* and *B* separately, then project the activity found across conditions for these tasks onto their corresponding PCs to obtain the latent activity matrices *L_A_* and *L_B_*. CCA begins with a QR decomposition of the transposed latent activity matrices as such 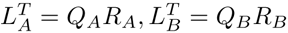. Singular value decomposition is then performed on the inner product matrix of *Q_A_* and *Q_B_* as such

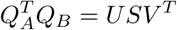

The elements of *S* contains the sorted canonical correlations which we use to determine the similarity between latents across tasks. We use the same pairs discussed in Section 4.3.

For Procrustes, we do not perform PCA on network activity before testing alignment. Procrustes optimizes the following objective

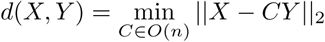

In this case, *X* and *Y* are *T* x*N* dimensional time-series, and *O*(*n*) is the orthogonal group *C^T^ C* = *I*. This metric will compare the similarity between the temporal patterns produced by each trajectory regardless of their orientation, offset, or scaling.

### 4.5 Targeted Dimensionality Reduction

To find evidence of explicit network primitives, we use targeted dimensionality reduction (TDR) to find axes in state space that explicitly encode task variables such as the frequency and rotational properties of different movements. Importantly, the existence of such axes in the network is purely emergent, as the one-hot rule input does not explicitly encode any task relationships. This is distinct from condition specific inputs, such as speed and direction, where input values increase along a single mode or move along a ring, respectively.

The TDR method relies on multi-variable linear regression for finding network axes. We first gather data from our model across all tasks and direction conditions and label them according to their specific frequency and rotational properties. Then, we cast the activity of units as a linear combination of task variables

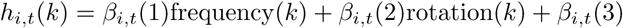

where *h_i,t_*(*k*) is the activity of unit *i* at time *t* for a trial *k*, frequency(*k*) is the frequency label for the movement, and rotation(*k*) is the rotation label for the movement. The frequency values include [0, 0.5, 1] for the reach, curved reach, and sinusoidal tasks respectively, and the rotation values are [1, 0, 1] for the curved reaches, reach, and sinusoid tasks respectively. Each *β_i,t_*(*v*) for *v* = 1, 2, 3 represent the coefficients that determine how much the trial-by-trial activity of unit *i* depend on each task variable.

To solve for the regression coefficients, we compute an *N*_coeff_ × *N*_trial_ matrix *F_i_* for each unit, where *N*_coeff_ is the number of coefficients (3), and *N*_trial_ is the number of trials used for that neuron. Note that since we can capture the activity of each unit for each condition in our model, *N*_trial_ is the same across units. The coefficients can be expressed as

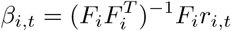

where each element *β_i,t_*(*v*) relates the task variable to unit activity.

After computing the coefficients *β_i,t_*, we then denoise them using PCA computed using network activity across tasks within a respective epoch, in the case of Figure 4 we focus on delay epoch activity. We then gather each vector *β_v,t_*(*i*), which is simply a transpose of the coefficient vector *β_i,t_*(*v*), and project it onto the first 12 components represented by the matrix *D*

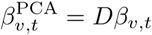

Next, we find the time 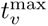 at which the vectors 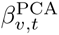 has maximum norm as such

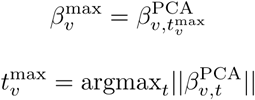

where each 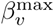 is a vector of length *N*_units_. Lastly, we orthogonalize each of the component directions using a QR decomposition

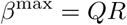

where *β*^max^ is a matrix where each column represents the component vectors 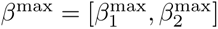, *Q* is an orthogonal matrix, and *R* is an upper triangular matrix. Here, *Q* represents the orthogonalized coefficient vectors, which we use to project activity onto to visualize differences in task representation. To visualize delay epoch representations along each of these axes, we project network activity at the last timestep of the delay epoch averaged across direction conditions onto *Q*.

### 4.6 Rule Input Alignment

Static rule inputs dictate the changes in network dynamics that utilize the proper motifs for a given task. The rule input vector, which is given by multiplying the one-hot rule input with the corresponding input weight, is a shift in a single direction in state space that allows for proper movement initialization and execution. As discussed in the main text, the rule input vector can either work to properly initialize the network state for movement, leaving movement related dynamics largely unchanged across tasks, or it could more dramatically shift movement-related dynamics leading to individualized solutions. To test these hypotheses, we create an alignment method between rule input vectors and a chosen subspace.

We compare rule input alignment with the delay and movement subspaces computed using PCA across all tasks and direction conditions. We use the first 5 and first 17 PCs of each subspace respectively, explaining over 90% of the variance for each respective epoch. Let *D* denote the subspace components (delay or movement), with *w* being the *N*_components_ (5 for delay and 17 for movement) dimensional vector containing the variance explained by each component. We take the rule input vector 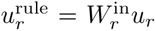, where *r* denotes the particular task chosen, and compute the absolute value of the inner product between 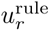 and each subspace component, then take the average weighted by their variance explained ratios

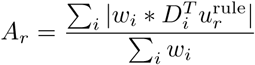

This metric is agnostic to whether the input is in the same or opposing direction to a particular component, and only tests the strength of alignment. Importantly, each PC component is of unit length, therefore strong alignment with a particular mode can only be produced if the rule vector has large magnitude in that direction. This metric also scales the importance of each mode to ensure that alignment is primarily along dominant modes.

### 4.7 Composite Rule Inputs

In order to test the model’s ability to combine primitive motifs to form novel movements, we tested whether or not motifs learned from other tasks can be uniquely combined in order to perform a separate task. Let *u^e^* ∈ ℝ^5^ denote the first five components of *u* for a particular condition. These components represent the rule inputs corresponding to the extension tasks. We optimize for a *ζ* ∈ ℝ^5^ element-wise scaled by *u^e^* such that 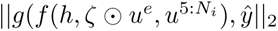, *ŷ*∥_2_ is minimized, where *N_i_* is the input size. For each desired task *ŷ*, the rule input corresponding to that task in *u^e^* is 0. We learn distinct rule inputs across conditions for better performance.

### 4.8 Sequencing Movement Patterns

To test if individual movement patterns can be sequenced, we switch the rule inputs corresponding to extension and retraction tasks at the middle of the movement epoch. Let 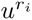 denote the input for task *i* and 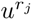 denote the rule input for task *j*. The total input to the network is then *u* = 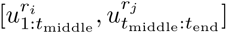, where *t*_middle_ and *t*_end_ denote the middle and end of the movement epoch respectively. In this setting, 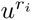 is always an extension task (e.g., Reach) and 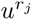 is always an extension-retraction task (e.g., ReachBack).

### 4.9 Fixed Point Analysis

In order to assess dynamical similarities across task pairs, we analyze the network fixed point structures and local dynamics around the fixed points. We specifically interpolate between the rule inputs for different tasks and determine whether the fixed point structure is preserved [9]. We perform this separately for different epochs of the motor task. To find fixed points, we follow methods used in [9, 59]. We optimize to find a set of hidden states 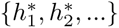 that minimize *F* (*h^∗^, u^∗^*), where *u^∗^* is a fixed input along the interpolated trajectory *αu*_1_ + (1 − *α*)*u*_2_, and *F* (*h, u*) is 1. In this case, *u*_1_ and *u*_2_ are the inputs for two different tasks, either at the end of the delay epoch or the middle of the movement epoch.

In addition to finding fixed points, we assess their local dynamics to determine if changes in stability occur as we interpolate. To do so, we linearize the network about the fixed point *h^∗^*:

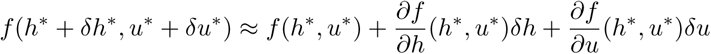

By definition, *f* (*h^∗^, u^∗^*) = 0, and second order terms are approximately zero given that ∥*δh*∥^2^ ≈ 0. Additionally, we keep our desired input *u^∗^* constant, therefore we can ignore any changes from *δu*. Thus, our Taylor expansion simplifies to

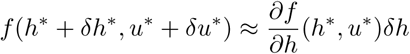

The eigenvalues of the linearized system’s Jacobian, 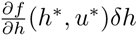, tells us the local dynamics near fixed points. As we interpolate from one task to another, we find the largest eigenvalue of the computed Jacobian for each fixed point found during optimization to assess changes in stability. To choose the fixed points used for plotting, we first select the fixed point closest to the state trajectory at *α* = 0. Afterwards, we choose the fixed point at *α_i_*_+1_ closest to the one previously selected for *α_i_*. We chose this method to best capture the trajectory of a single fixed point as we interpolate from one task to another. In addition to viewing max eigenvalues, we additionally quantify change in the structure of fixed points by taking their euclidean distance, 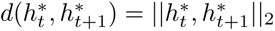, during interpolation. This metric uses the same method for selecting fixed points as previously mentioned.

### 4.10 Dynamical Similarity Analysis

In Section 2.3 we perform DSA to identify clusters of dynamical and geometrical similarity between tasks. Given two dynamical systems

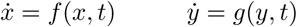

DSA samples data from each system, 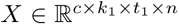 and 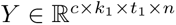, where *c* is the number of conditions, *k* is the number of trials per condition, *t* is the number of timepoints, and *n* is the number of units in the system. Then, the data is used to create delay-embedded Hankel tensors with lag of 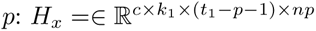 [60]. The Hankel Alternative View of Koopman (HAVOK) approach is then used to build the necessary dynamic mode decomposition (DMD) matrices. This approach flattens all but the last dimensions of *H* and fits reduced rank regression models with rank *r* to the coordinates of the next time step 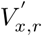:

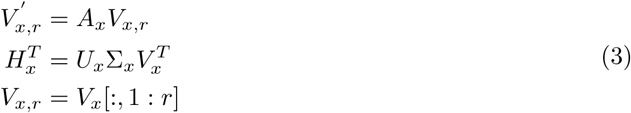

The resulting DMD matrices are given by *A_x_* and *A_y_*. Lastly, Procrustes analysis is performed to test the alignment between the DMD matrices through a similarity transformation. The optimization problem posed by Procrustes analysis over vector fields is given as:

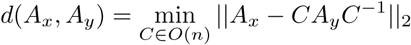

In Figure 3(g) we perform this analysis on separate tasks to test their dynamical similarity. Instead of using the ten tasks in totality, we also split the extension and retraction portions of the extension-retraction tasks to more clearly test the learned dynamical structure across movement patterns. To test the strength of the clustering given by the labeled extension and retraction tasks, we take the silhouette score of each cluster, defined as

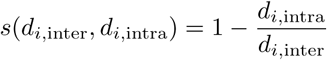

Where *d_i,_*_inter_ is the average distance between unit *i* and every unit in the nearest cluster, and *d_i,_*_intra_ is the average distance between unit *i* and every unit in the same cluster. The average across all units *i* is used to score a cluster. A low score denotes weak clustering, while a score close to one denotes otherwise.

### 4.11 Transfer Learning

To test whether the network can reconfigure learned motifs to quickly learn new tasks, we train the network on all tasks except CClkCR and CClkCycle, then freeze the learned weights *W_h_, W*_out_, *W*_in_, *b_h_, b*_out_ and only learn two new columns for the transfer input weights 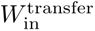. We exclude both CClkCR and CClkCycle to ensure that the network does not learn any counter-clockwise curved reach motif during any task. To test whether the motifs learned during pretraining are necessary for performing the transfer learning tasks well, we train the model on stricter subsets of pre-trained tasks to determine if performance on the held out tasks are affected. The additional task subsets are as follows:

- sin: {Sinusoid, InvSinusoid, Figure8, InvFigure8}
- reach: {Reach, ReachBack}
- nosin: {Reach, ClkCR, ReachBack, ClkCycle}
- nocr: {Reach, Sinusoid, InvSinusoid, ReachBack, Figure8, InvFigure8}
- cr: {ClkCR, ClkCycle}

We additionally test whether learned subspace structures are reused during transfer learning. To do so, we compute the angular distance and disparity between CClkCR with all other extension and retraction movements, and similarly for CClkCycle.

### 4.12 Varying Hyperparameters

We trained networks with variable activation functions and hidden units to test if the subspace structures found using the baseline model are consistent across hyperparameters. The activation functions include Softplus

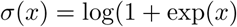

the ReLU function

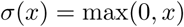

and the Tanh function

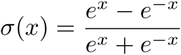

We test models with 128, 256, and 512 hidden units for each activation function. Supplemental Figure 3 includes the results of this hyperparameter search, showing that shared low-dimensional subspaces across tasks are consistently used in different settings.

### 4.13 Task Variance Analysis

In Supplemental Figure 1, we perform a task variance analysis to determine if populations of neurons are differentially selective across tasks. We use the same task variance metric as [8], where the variance is computed across all conditions and timesteps for a particular task then averaged. The form of the task variance metric is given as

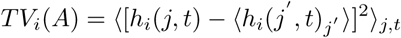

where *i* denoted the current unit, *A* represents the task, *j* is the condition, and *t* is time. In our case, we compute this metric over all speed and direction conditions during the movement and delay epochs. Units that have a summed task variance across all tasks less than 10^−3^ were discarded for inactivity.

Given the task variance of each unit, we aim to find clusters of task selectivity in the population. To do so, we used k-means clustering on the task variances while ranging the number of clusters from 2 to 20. To choose the best clustering, we compute the silhouette score, averaging over all units, then choose the highest score.

