## Supplemental Figures for "Multitasking Recurrent Networks Utilize Compositional Strategies for Control of Movement"

### Supplemental Material

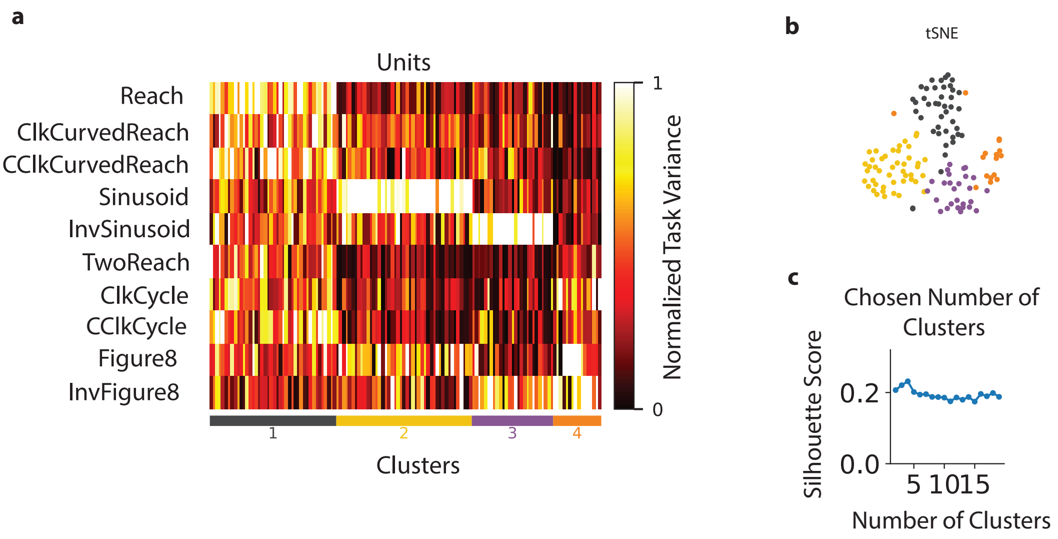


**Fig. 1** Network modularity across tasks. (a) Normalized task variance of each unit for each task, with the corresponding clusters of units identified. The model contains a total of four clusters across ten tasks. (b) Visualization of the task variance clustering using tSNE. (c) Silhouette score as the number of clusters for k-means clustering increases. The final number of clusters is chosen as the highest score.

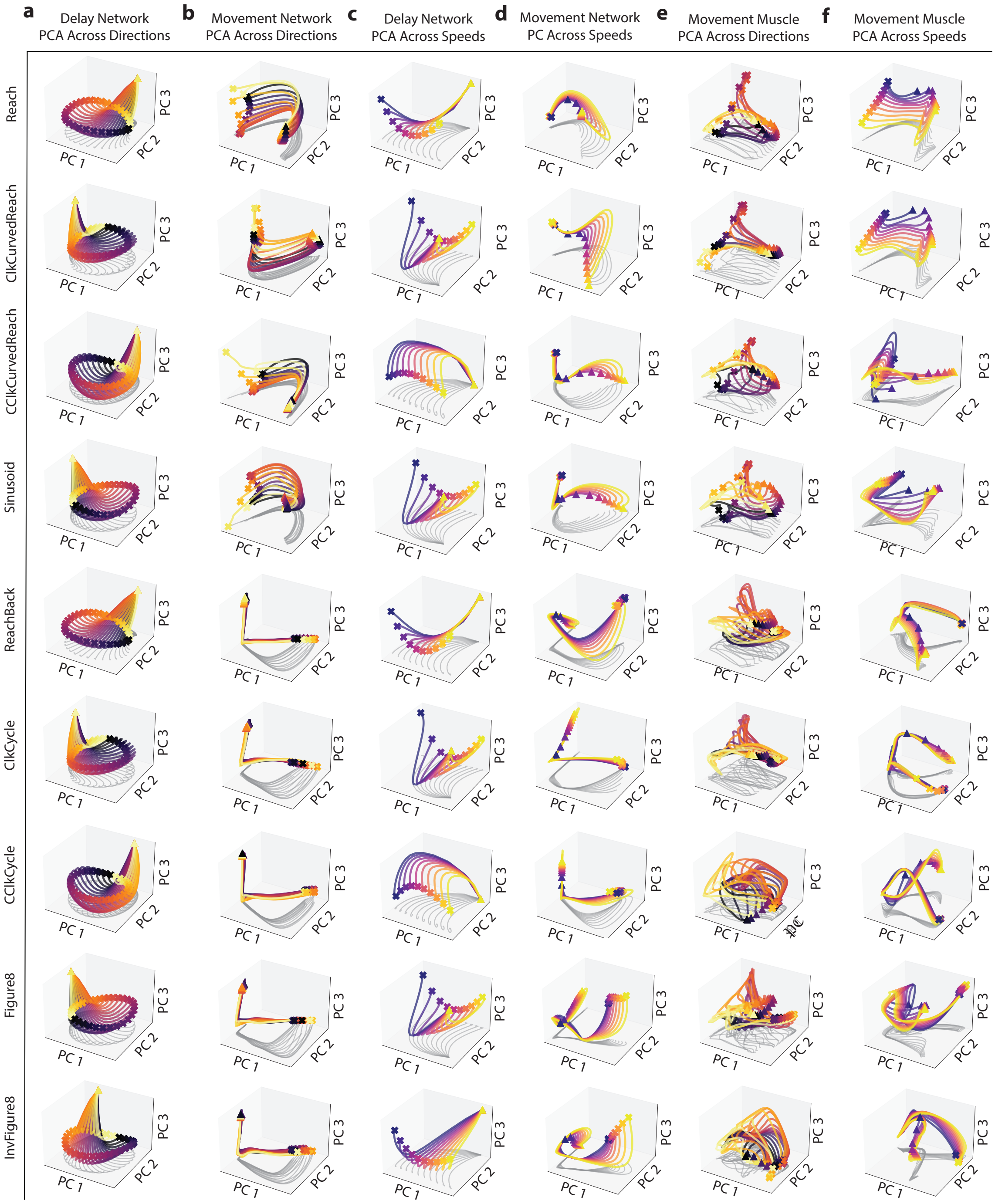


**Fig. 2** All delay and movement PCs for network and muscles. (a) Network trajectories projected onto top three PCs found during the delay across direction conditions for nine tasks (not including InvSinusoid). Each task is shown in different rows. (b) Same as (a) but using movement activity for PCA and projection. (c) Same as (a) but across speed conditions. (d) Same as (b) but across speed conditions. (e) Same as (b) but in the muscle space. (f) Same as (d) but in the muscle space. All tasks use similar strategies to the InvSinusoid task displayed in the main text.

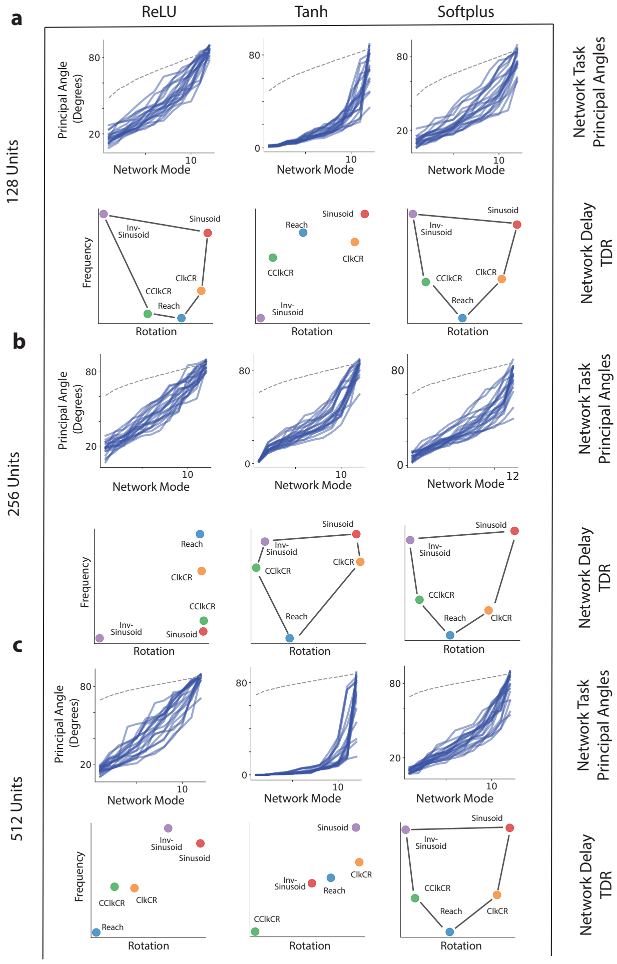


**Fig. 3 Shared subspaces are robust to network hyperparameters; structured task geometry is not** (a) (Top row) principal angles across similar tasks for networks with different activation functions and 128 units. (bottom row) TDR during the delay epoch across networks with varying activation functions and 128 units. (b) Same as (a) but for networks with 256 units. (c) Same and (a) and (b) but for networks with 512 units. Softplus networks consistently find structured rotation and frequency axes, but this is not always present across other activation functions. Low dimensional subspaces however are consistently utilized across activation function and varying unit numbers.

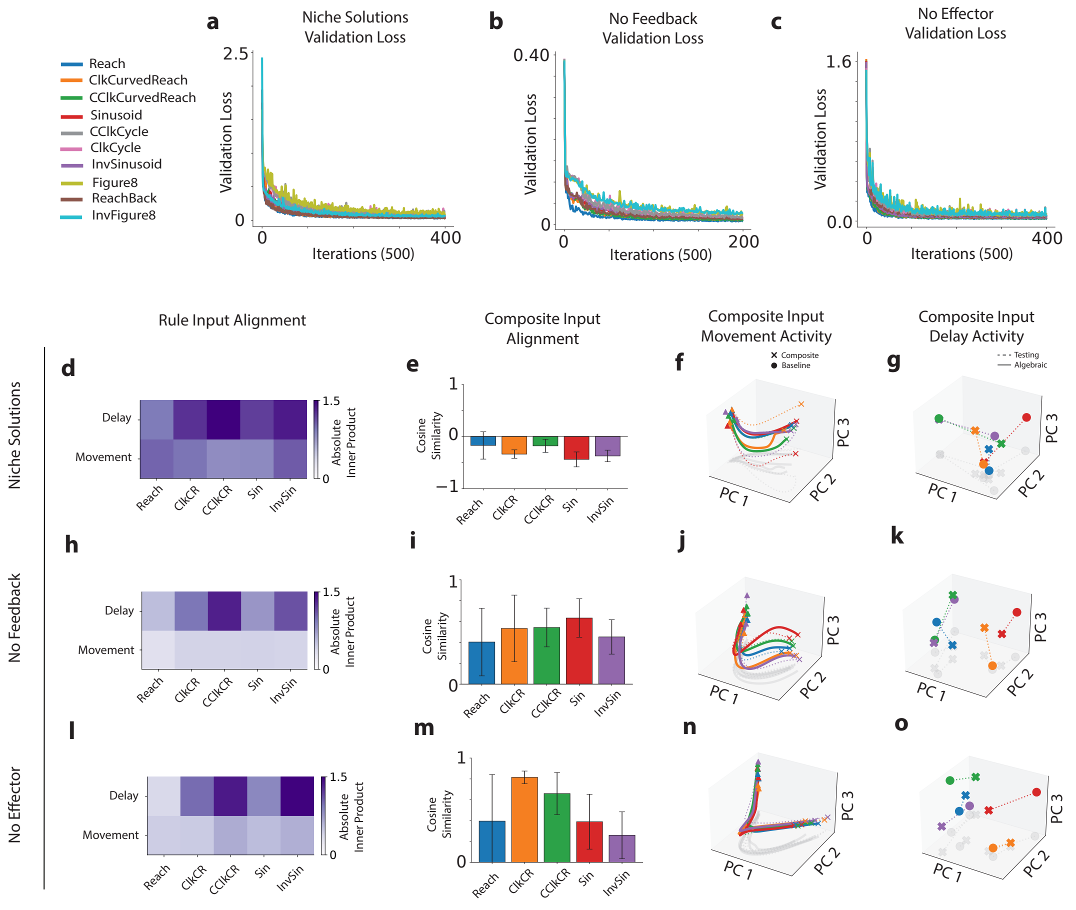


**Fig. 4 Compositional strategies of ablation models.** (a) Validation loss for Niche Solutions model. (b-c) Same as (a) for No Feedback and No Effector models respectively. Each network trains well across tasks until the loss plateaus. (d) Rule input vector alignment for the Niche Solutions model with the delay and movement subspaces across tasks. A weighted average is found for the absolute value of the inner product between each subspaces' components and the rule input weights for a particular task, weighted by the variance explained by an individual component. (e) Cosine similarity between the standard and composite rule input weights for the Niche Solutions model, reduced to the first 18 PCs found during all epochs and conditions across tasks. Rule vectors are less aligned than the baseline model. (f) Movement activity using the standard and composite input, movement PCs found using standard rule input. (g) State of the network at the end of the delay epoch for the standard and composite rule input, with PCs captured using the standard rule input. Initial conditions are less aligned than the baseline network. (h-k) Same as (d-g) but for the No Feedback model. (l-o) Same as (d-g) for the No Effector model.
